# A framework to infer *de novo* exonic variants when parental genotypes are missing enhances association studies of autism

**DOI:** 10.1101/2025.07.24.666675

**Authors:** Haeun Moon, Laura Sloofman, Marina Natividad Avila, Lambertus Klei, Bernie Devlin, Joseph D. Buxbaum, Kathryn Roeder

**Affiliations:** Department of Statistics, Seoul National University, Seoul, South Korea; School of Transdisciplinary Innovations, Seoul National University, Seoul, South Korea; Institute for Data Innovation in Science, Seoul National University, Seoul, South Korea; Seaver Autism Center for Research and Treatment, Icahn School of Medicine at Mount Sinai, New York, New York, USA; Department of Psychiatry, Icahn School of Medicine at Mount Sinai, New York, New York, USA; Department of Genetics and Genomic Sciences, Icahn School of Medicine at Mount Sinai, New York, New York, USA; Friedman Brain Institute, Icahn School of Medicine at Mount Sinai, New York, New York, USA; Department of Neuroscience, Icahn School of Medicine at Mount Sinai, New York, New York, USA; The Mindich Child Health and Development Institute, Icahn School of Medicine at Mount Sinai, New York, New York, USA; Department of Psychiatry, University of Pittsburgh, Pittsburgh, Pennsylvania, USA; Department of Statistics, Carnegie Mellon University, Pittsburgh, Pennsylvania, USA; Computational Biology Department, Carnegie Mellon University, Pittsburgh, Pennsylvania, USA

**Keywords:** *de novo* mutation, classification, genetic association, autism spectrum disorder, congenital heart disease

## Abstract

**Motivation:** Gene-damaging mutations are highly informative for studies seeking to discover genes underlying developmental disorders. Traditionally, these *de novo* variants are recognized by evaluating high-quality DNA sequence from affected offspring and parents. However, when parental sequence is unavailable, methods are required to infer *de novo* status and use this inference for association studies.

**Results:** We use data from autism spectrum disorder to illustrate and evaluate methods. Separating *de novo* from rare inherited variants is challenging because the latter are far more common. Using a classifier for unbalanced data and variants of known inheritance class, we build an inheritance model and then a *de novo* score for variants when parental data are missing. Next, we propose a new Random Draw (RD) model to use this score for gene discovery. Built into an existing inferential framework, RD produces a more powerful gene-based association test and controls the false discovery rate.

**Availability and Implementation:** The implementation code and publicly available data are provided at: https://github.com/HaeunM/TADA-RD.

## Introduction

Rare genetic variation has been key in the identification of specific genes that alter development, including genes underlying congenital heart disease (Jin et al., 2017; Watkins et al., 2019) and autism spectrum disorder (Fu et al., 2022). For ASD, most of the information inherent in rare variation comes from *de novo* exonic mutations (Fu et al., 2022), and the same is true for chromatin modifiers underlying some forms of congenital heart disease (Jin et al., 2017). This presents challenges for study design. To identify a variant as *de novo*, definitively, requires high-quality whole-exome or whole-genome sequence data from an affected offspring (the proband) and both parents. An alternative design contrasts probands with appropriately matched controls. While the case-control design contributes some evidence for association, the information gleaned from rare variants in such a cohort is typically modest (Fu et al., 2022), in large part because which variants are *de novo* and which are inherited is hidden. We propose methods to recover this information.

This problem is challenging because there are far more inherited than *de novo* variants per subject. Consider autism spectrum disorder (ASD) as an example. Defining a rare variant as having a minor allele frequency *MAF <* 0.001, a typical person diagnosed with ASD carries one to two *de novo* variants in their exons, compared to approximately thirty-fold more rare inherited variants. All classification problems are challenged in this setting. In addition, the classification of harmful from benign variants remains error-prone, although *de novo* variants that affect liability must alter gene function and neurodevelopment. Furthermore, because only about 5% of the coding genes have a strong impact on ASD, and many of them are not yet known, not all *de novo* variation matters. Any model to infer *de novo* variation will benefit from information about association for specific genes.

For the problem of sorting *de novo* from inherited variants, frequencies of variants carry information. If a putative *de novo* variant is relatively common in the population, it is more likely to be an inherited variant that went undetected in the parents due to randomness of the sequencing process. Using this logic, we can reasonably expect that most true *de novo* variants are rare in the sample under study and rare or absent in other population databases. In addition, genes in which functional variation is subjected to evolutionary negative selection, constrained genes, are important for typical development, ASD and many other developmental disorders. A useful measure of gene constraint is LOEUF. Genes with low LOEUF scores are constrained (Karczewski et al., 2020): in a population, they tend to have relatively few variants leading to loss-of-function of the gene. These loss-of-function variants, often annotated as protein-truncating variants or PTVs, are not the only kind of variation that can cause loss-of-function. However, in this work, we will focus on PTVs because they make up the majority of damaging variants found in ASD subjects, developing a classifier to separate *de novo* from inherited PTVs.

Next, we develop an approach to incorporate this information into a gene-based association framework and thereby garner greater power to identify relevant genes. We build on the TADA model (Transmitted and De novo Association) (He et al., 2013), which combines different types of data, including those of families with affected probands, probands without parental information and controls. Although TADA provides a comprehensive analytical framework to accumulate the signals from each kind of data and over types of variants (e.g., PTV, missense), it provides only a modest signal for association from case versus control data. If we can identify the *de novo* variants in cases and controls, we conjecture that this will enhance the TADA signal and the power for association. In this manuscript, we propose a new integrated model that evaluates variation in case and control samples, attempts to sort *de novo* mutations from inherited variation, and incorporates this information into a refined TADA model (Figure 1). We exemplify the method using published ASD data. Importantly, the approach we develop can be used to associate rare variants with any developmental phenotype, such as schizophrenia, obsessive-compulsive disorder, and congenital heart disease.

**Fig. 1.**
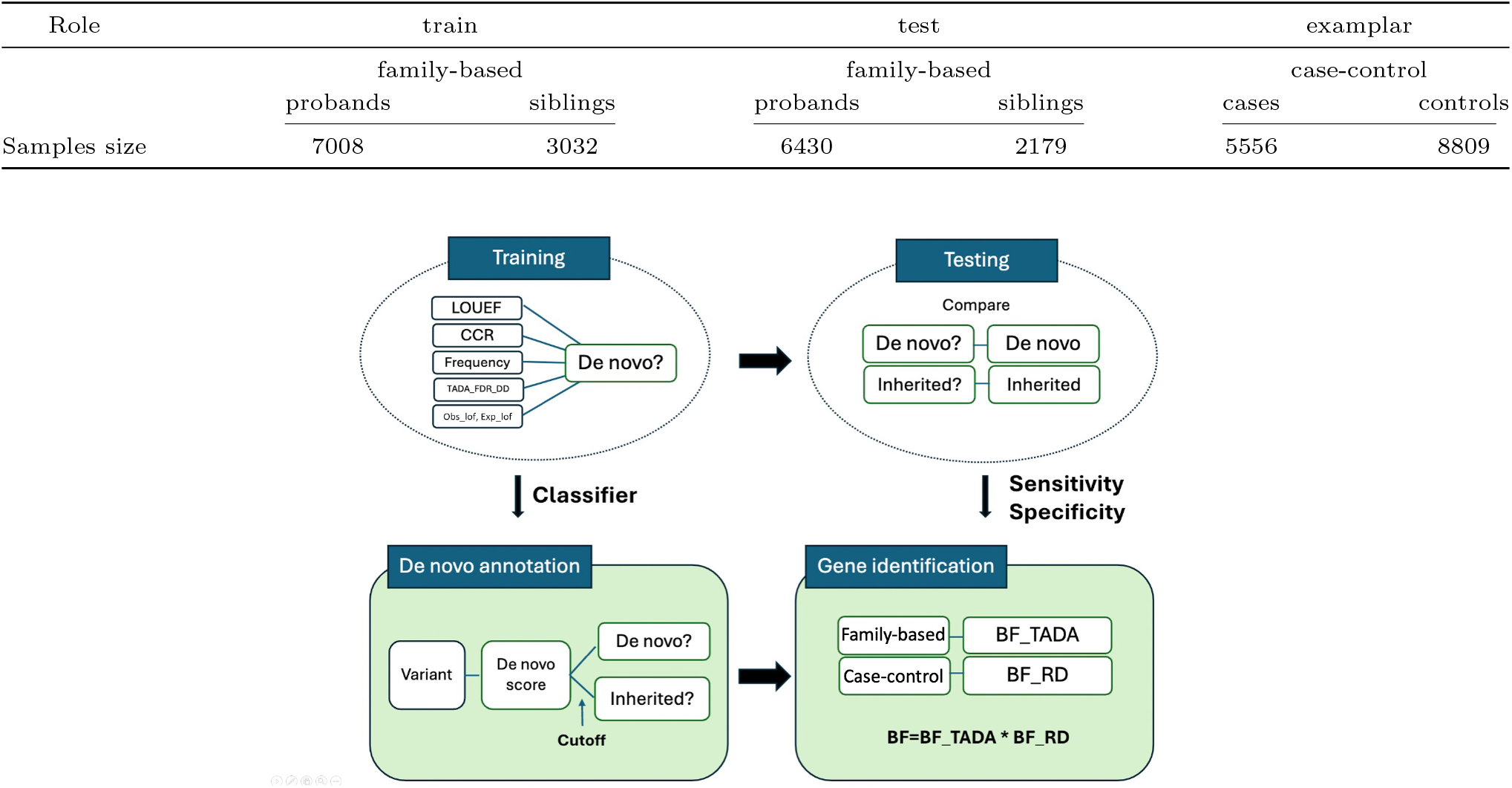
Overview of workflow and the datasets. **(Top)** Summary of dataset used in the data application. The training data is from SPARK. The family-based test data is from the ASC, including the Simon’s Simplex Collection, and the case-control data is from the ASC (Fu et al., 2022) **(Bottom)** Summary of the method. In pipeline 1, an unbalanced classification algorithm *ClassDn* is trained with variants of a known inheritance class utilizing offspring-level variant information (**training**) and then is tested on another set of variants with known inheritance class to learn parameters about the classification performance (**testing**). In pipeline 2, *ClassDn* is applied to variants without parental genotype information to learn their *inferred de novo* status (**de novo annotation**), and *Random Draw model* is applied to them to obtain BF RD for each gene. Bayes factors obtained by multiplying BF RD by BF TADA from variants with known inheritance classes represent the association score of genes (**gene identification**)

## Materials and Methods

### Overview of workflow and the datasets

Our procedure comprises two separate pipelines (Figure 1). The first builds *ClassDn*, a classifier to predict the inheritance class of variants (*de novo* or inherited) using offspring level variants with known inheritance class. *ClassDn* is then applied to a test dataset to assess the classifier’s performance, which serves as a component of the input parameters for the data application. In the second pipeline, *ClassDn*’s predictions are applied to variants without parental information to obtain their likelihood of being *de novo*, called de novo scores. When applied to the full case sample, these scores are converted to gene-level risk scores using a newly proposed procedure called the *Random Draw (RD)* model, which is designed to be robust to uncertainty in the predicted inheritance class and to control gene-level false discoveries. Combining this information with other information assimilated in the existing TADA framework (Fu et al., 2022) produces a more powerful gene-based association test.

Data from previously published exome sequencing data from ASD and control subjects are used for our analysis. These data, described in Fu et al. (2022), are derived from two sources: the Simons Powering Autism Research (SPARK) initiative and the Autism Sequencing Consortium (ASC) samples. In this study, we only focused on putative protein-truncating variants or PTV, because they are best understood in terms of their impact on gene function. Following previous analyses by the ASC consortium, we restrict our analysis to ultra-rare variants, defined as those present at *<* .1% population frequency. Both the ASC and SPARK datasets consist of two types of data: family-based data, where the inheritance class of offspring variants is known, and case-control data, where such information is not available. For the first pipeline of our method, SPARK family-based data are used to train *ClassDn* using unbalanced classification algorithms, and the ASC family-based data are then used to learn the performance parameters of *ClassDn*. For the second pipeline, ASC family and case data are utilized to obtain integrated association scores that constitute risk gene test statistics (see Figure 1).

### Classification algorithm for inheritance class

A major source of signal for the association of genes with ASD arises from *de novo* PTVs; by comparison, the information derived from the case-control and the inherited variants is relatively minor (Fu et al., 2022). Moreover, the signal derived from case-control samples is far greater than that from inherited variation assessed from family-based samples (Satterstrom et al., 2020), indicating that the case-control sample likely includes a non-negligible number of *de novo* and very recent mutations, which are typically more damaging. We conjecture that the distinct nature of *de novo* and inherited variants will allow us to separate them using only offspring genetic information, which includes both variant-and gene-level variables. Here we use family-based data to build the *ClassDn* classifier to identify *inheritance class*.

We incorporate six variables to infer the inheritance classes of variants: Allele Frequency (AF), LOEUF, CCR, FDR TADA DD, obs lof and exp lof. Allele Frequency refers to the proportion of a particular allele (variant) within a population, which is obtained from the gnomAD library. Gene (LOEUF) and variant (CCR) scores of the evolutionary constraint are also incorporated. A “strong” constraint score indicates that changes in the region or gene are likely to reduce the reproductive success of individuals. Consequently, PTVs falling in these regions/genes are more likely to be *de novo* than inherited. A variant-level constraint score, a constrained coding region (CCR), measures how depleted specific areas within coding regions are in protein-changing variants (Havrilla et al., 2019); it ranges from 0 to 100. CCR scores are matched to variant locations using starting position of the CCR. A gene-level score, LOEUF is a continuous measure of selective pressure against loss of function of a copy of a gene (Karczewski et al., 2020): obs_lof and exp_lof are two fundamental components of the LOEUF score and individually contribute to distinguishing *de novo* variants in classifiers. Because ASD is often one of a constellation of symptoms of neurodevelopmental delay (NDD) or developmental delay (DD/NDD), genes affecting ASD and those affecting DD/NDD largely overlap (Doshi-Velez et al., 2014; Sanders et al., 2019). Therefore, we use the known risk score of gene associated to one of theses conditions TADA_DD_FDR (Fu et al., 2022). All six covariates effectively distinguish the inheritance class of variants (Figure 2A). Compared to inherited variants, *de novo* variants exhibit a lower allele frequency, a lower LOEUF score, a higher CCR and a lower FDR score related to DD. Such distinction confirms that a classifier based on these variables will generate meaningful separation of variants in terms of different inheritance classes.

**Fig. 2.**
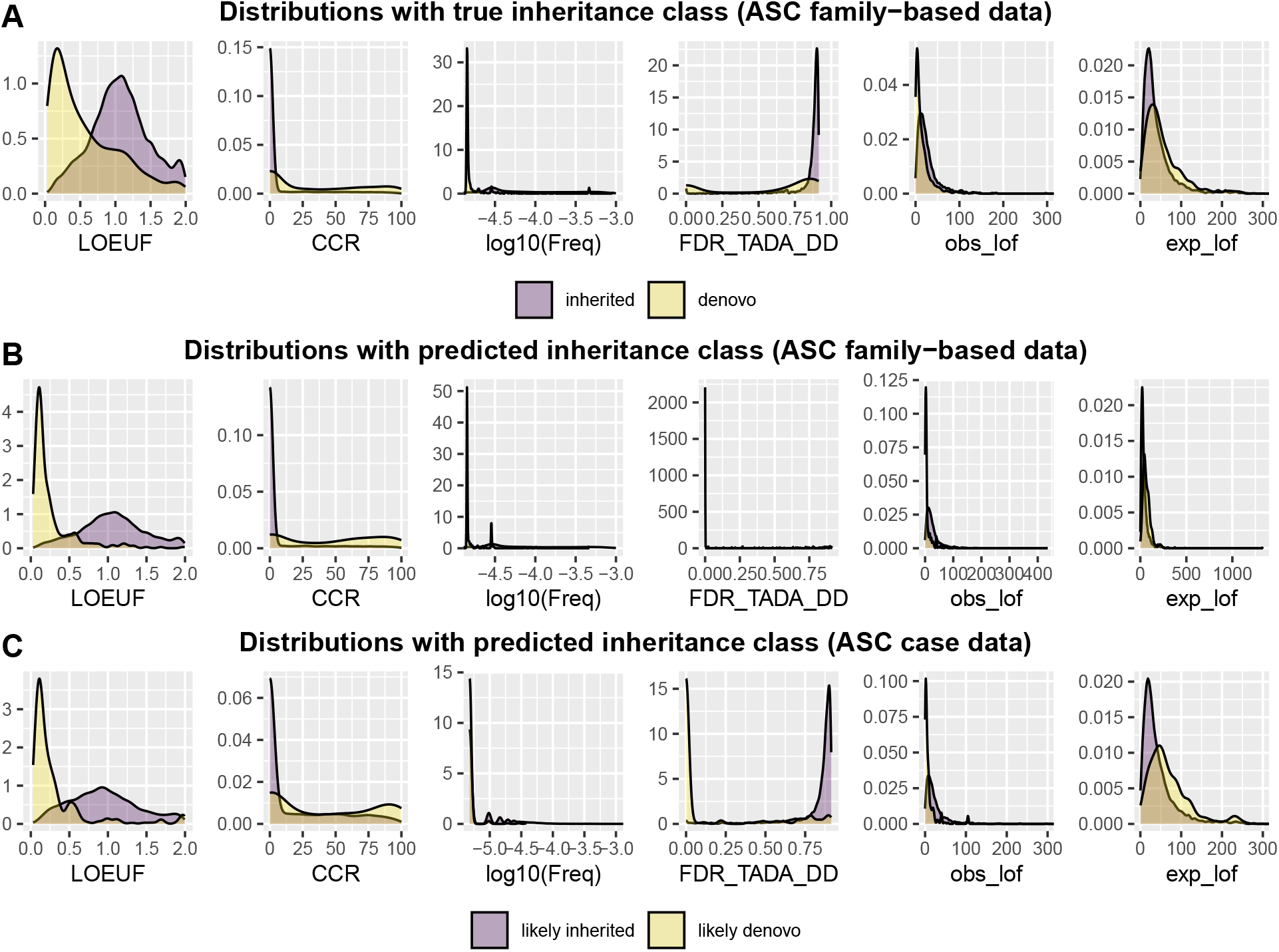
Six variables used in the analysis help distinguish *de novo* and inherited variants. **A**. Density plots of covariates for *de novo* and inherited variants in proband data from the ASC family-based data. **B**. Density plots of covariates for likely *de novo* and likely inherited variants in proband data from ASC family-based data. **C**. Density plots of covariates for likely *de novo* and likely inherited type of variants in ASC case data. RUSboost with a threshold 0.7 was utilized to generate the Figure 2B and 2C.

Notably, the number of inherited variants greatly exceeds that of *de novo* variants. In such imbalanced data, a naive use of classification algorithms can lead to neglecting the performance of minority class data. Therefore, for *ClassDn*, we implemented algorithms specifically designed to handle imbalanced data, such as RUSBoost (Seiffert et al., 2009) and Underbagging (Barandela et al., 2003), both of which balance class prediction through resampling techniques. The algorithms output the likelihood of being *de novo* for each variant, termed de novo scores. Combining this with a specific choice of threshold, we classify a variant as “likely de novo” if its de novo score exceeds the threshold and “likely inherited” otherwise. The overall performance of *ClassDn* depends on the choice of threshold and can be summarized with two parameters: sensitivity (the probability that *de novo* variants are classified as *de novo*; *w*_1_) and specificity (the probability that inherited variants are classified as inherited; *w*_2_). Naturally, a trade-off exists between sensitivity and specificity: using a higher threshold decreases sensitivity and increases specificity. The overall performance in our application is summarized in Figure 3; for RUSboost with a choice of threshold 0.7, the sensitivity is 0.164 and the specificity is 0.998 and for Underbagging, the sensitivity is 0.335 and the specificity is 0.990. The separation between two predicted classes is notable with six covariates (Figure 2B, RUSBoost, with threshold 0.7, was used to generate this figure). This algorithm, being conservative in predicting the *de novo* class, results in a slightly more extreme separation compared to the true inheritance class for some covariates.

**Fig. 3.**
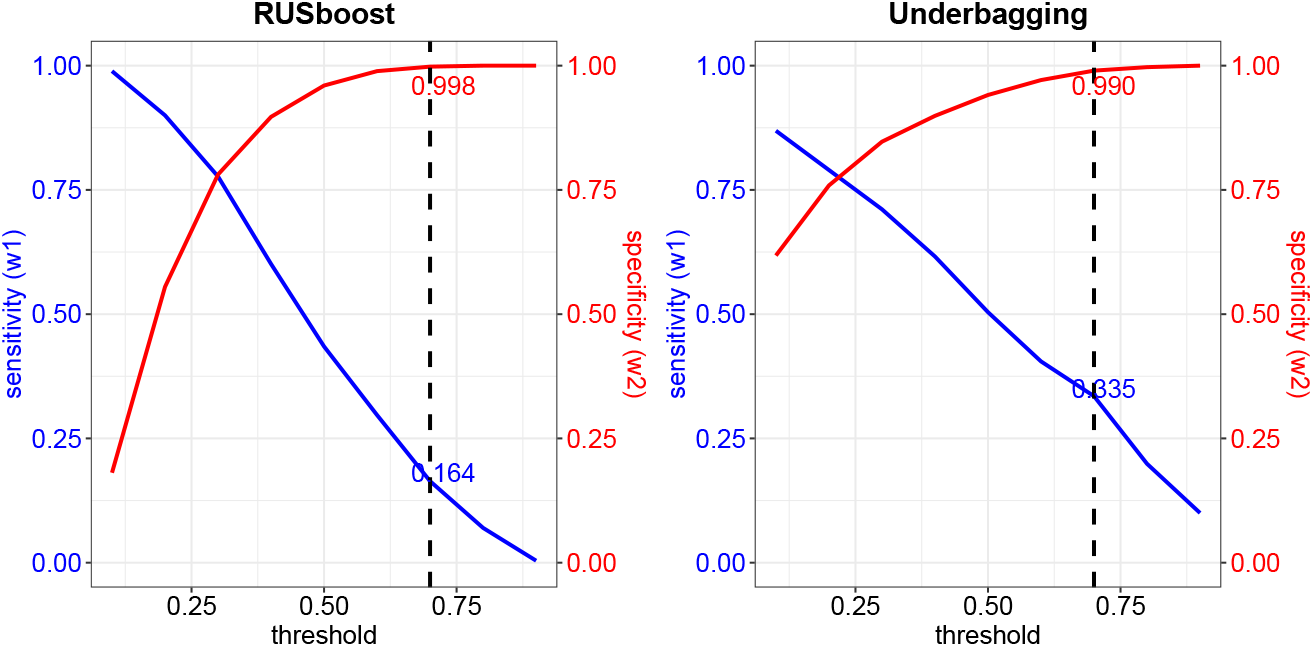
Sensitivity and Specificity of classifiers for a range of thresholds. RUSboost (left) and Underbagging (right). Results at the threshold of 0.7 are indicated by the dashed line.

### Identification of risk genes

The Transmission and De Novo Association (TADA) model is a Bayesian statistical framework to identify associated genes using variants of various inheritance classes (He et al., 2013) of family-based and case-control data. The original model was refined in Fu et al. (2022) to include a wider variety of genetic variation, including multiple classes of PTV, damaging missense variants, and copy number variants. Here we advance the TADA model by using *ClassDn* to interpret PTV variants found in individuals with ASD, but for whom parental data are missing (case-only data). This information is summarized as an association score (Bayes Factor) using the *Random Draw* model, and it replaces the previous component of the TADA model involving a case-control contrast of distributions. The final TADA model, then, integrates scores from all sources of information to discover associated genes.

The Random Draw model assumes that each variant is a random draw with a given probability of being *de novo*. This probability is assumed to depend on the risk status of the gene, with risk genes expected to have a greater fraction of *de novo* variants. Consequently, a gene with a higher proportion of *de novo* variants should produce a higher risk score, a trend that is strongly supported with data (Suppl Figure S2). Once we assume some prior distribution for the probability of drawing a *de novo* variant under each risk and non-risk gene scenario, we can compute the likelihood for the observed count of *de novo* and inherited variants and then contrast the two scenarios using a Bayes factor. A remaining challenge for implementation is that the predicted inheritance class from *ClassDn* is only a proxy for the real inheritance status of variants. To incorporate such uncertainty, classification accuracy parameters are utilized.

Define *I* to be an indicator of *de novo* status of a given variant (1 for *de novo* and 0 for inherited) and *X* to be its predicted inheritance class (1 if the ClassDn score is greater than the threshold, and 0 otherwise). We denote the classification accuracy parameters as *w*_1_ = *P* (*X* = 1|*I* = 1) and *w*_2_ = *P* (*X* = 0|*I* = 0). For an indicator *D* of the risk status of the gene (1 for risk genes and 0 for non-risk genes), and a mutation rate *µ*, define 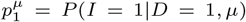 and 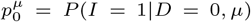 to be probability that the variant is *de novo* under each scenario 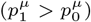. Suppose we observe counts of (*x*_*d*_, *x*_*h*_) variants for a given gene, where *x*_*d*_ represents the number of likely de novo variants and *x*_*h*_ represents the number of likely inherited variants, classified as greater than and less than the threshold, respectively. The likelihood of such data under each scenario are identified from the binomial distributions, as

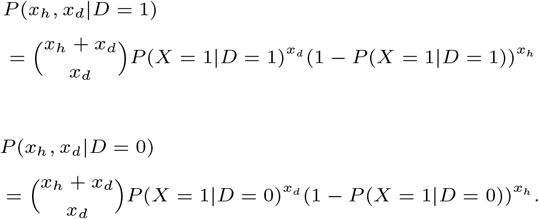

For given values of 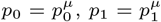, the binomial parameters can be expressed as *P* (*X* = 1|*D* = 1) = *w*_1_*p*_1_ +(1 − *w*_2_)(1 − *p*_1_), and *P* (*X* = 1|*D* = 0) = *w*_1_*p*_0_ + (1 − *w*_2_)(1 − *p*_0_). Finally, the evidence for *D* = 1 versus *D* = 0 can be calculated as a Bayes factor, which accounts for uncertainty in the parameters (*p*_0_, *p*_1_) using prior distributions *P*_0_ and *P*_1_. For details, see the Supplementary Material A.

## Results

For illustration, we restrict the data input of the TADA model to two classes of variants: *de novo* PTVs obtained from family data and PTVs from case and control samples. Our objective is to evaluate the performance of the Random Draw model, relative to the usual treatment of case-control variants in the TADA model. Toward this end, the TADA statistic is obtained as a product of the BFs obtained from *de novo* PTVs and the Random Draw model (TADA_*RD*_) or case-control model (TADA_*CC*_).

### Validation of the Random Draw Model

The Random Draw model provides a new way to identify risk scores from variants without parental information. It shares similarities with TADA_*CC*_ model in the sense that both assign moderate to large scores to genes with more *de novo* variants, which helps in the identification of risk genes. However, these two models have a different perspective in how to use unlabled variants obtained from case-control samples: the Random Draw model contrasts the count of inherited to *de novo* variants in cases only, and each additional inherited variant decreases the score; but the TADA_*CC*_ model compares the counts of unlabled variants in cases to controls. The difference arises from a fundamental choice for how to assess risk: the original TADA model assumes that an excess of either *de novo* or rare inherited variants is evidence of association; while the random draw model relies on the observation that risk genes have a much higher rate of *de novo* variants compared to non-risk genes. Moreover, the observed negative relationship between the number of inherited variants per gene and the final risk score of the genes supports the foundation of the Random Draw model (see Supp Figure S2).

Even though the Random Draw model is applied only to case-control data in real applications, as a check for the validity of the new model, we compare the risk scores obtained from the original TADA model and the Random Draw model using only the family-based dataset (with *w*_1_ = *w*_2_ = 1 for the Random Draw model). Despite the differences in formulation the resulting risk scores show high consistency (Supp Figure S3).

### Simulation

We conduct null and power simulations to verify that the proposed method enhances gene discovery while reasonably controlling false discoveries.

In the null simulation, we sample variants from the ASC unaffected sibling dataset, which includes complete covariate information, to generate a fully null case dataset with no ASD signal; the detailed procedure is described in the Supplementary Material B. For each null simulated dataset, we apply our *ClassDn* method to obtain de novo scores for each variant, identify the number of likely de novo and inherited variants for each gene with a choice of threshold *c*, and then apply the Random Draw model.

Obtained Bayes factors (BFs) from this dataset are multiplied by the BFs from the ASC family-based dataset. Final BFs are jointly transformed to FDR q-values following the approach described in Li et al. (2010) (Section 2.3). Two different types of cutoffs are used for gene selection; first, we convert q-values to p-values and selected genes with p-values less than 0.05 divided by the total number of genes considered—that is, using the Bonferroni procedure; second, we simply use a FDR cutoff of 0.05. For evaluation, the number of false selections (Per-family error rate: PFER) averaged over the datasets are reported. While we cannot know the ground truth, previously published results from a much larger sample and more genetic information are used to determine the set of ‘true’ genes to calculate the PFER (Fu et al., 2022). The benchmark method for comparison is the TADA model applied only to *de novo* PTVs in the ASC family-based dataset, where there is no prediction uncertainty. In this case, the PFER is 0 when using the Bonferroni cutoff and 8 when using a fixed cutoff of 0.05. In comparison, the Random Draw model with inferred *de novo* variants does not particularly inflate false discoveries, especially with thresholds of 0.7 or 0.9 (Table 1).

**Table 1.** A random draw approach enhances power while maintaining reasonable control of false discoveries in simulation datasets.

| c | Null (PFER) |  |  |  | Power |  |  |
| --- | --- | --- | --- | --- | --- | --- | --- |
| | RUSboost | | Underbagging | | $w_1$ | $w_2$ | Disc |
|  | Bonf | qval | Bonf | qval |  |  |  |
| 0.3 | 1.1 | 11 | 1.2 | 10.6 | 0.829 | 0.624 | 66.2 |
| 0.5 | 1 | 10.5 | 1.1 | 10.2 | 0.624 | 0.829 | 72.1 |
| 0.7 | 0.1 | 8.2 | 0.1 | 8.5 | 0.376 | 0.943 | 72.4 |
| 0.9 | 0 | 8 | 0 | 8 | 0.171 | 0.987 | 67.0 |

When only family-based data is considered, PFER is 0 with a bonferroni cutoff and 8 with the fixed cutoff of 0.05. The number of discovery is 58 when only family-based data is used, and 64 when case-control data is incorporated with TADA<sub>CC</sub> model.

In the power simulation, we use the list of genes and their corresponding number of variants from the ASC case dataset and simulate the inheritance class and de novo score of each variant; in reality this is hidden information in these data. Specifically, we assume that the fraction of *de novo* variants is 0.603 for genes with q-value *<*0.05 in the most informative data analysis published (Fu et al., 2022) and 0.026 otherwise. These values represent the proportion of *de novo* variants within the ASC family-based data for the risk gene and non-risk genes, respectively. Then the final numbers of *de novo* variants are computed as the product of these *de novo* fractions and the number of case variants of each gene, rounded to the nearest integer, and lower bounded by zero. The remaining variants are set as inherited variants. Based on the simulated inheritance class label for each variant, we assign a random de novo score generated from a Gaussian distribution, with a mean of 0.6 and a variance of 0.1 for *de novo* variants and a mean of 0.2 and a variance of 0.1 for inherited variants. In our analysis, we assume that the inheritance status of variants is unknown, but their de novo scores obtained from *ClassDn* are observable. For convenience, we further assume that the distribution of de novo scores is known in both *de novo* and inherited scenarios. This implies that once a threshold *c* is selected, the sensitivity and specificity parameters *w*_1_ and *w*_2_ can be computed from these distributions. In real-world applications, these parameters would need to be estimated using test data labeled with the true *de novo* status.

Upon generating the simulated data, we select a threshold level between 0.3 and 0.9 and apply it to the de novo scores: variants with scores exceeding the threshold are classified as likely de novo, while those below are classified as likely inherited. Given the sensitivity (*w*_1_) and specificity (*w*_2_) corresponding to the selected threshold, and the number of likely de novo and inherited variants per gene, Bayes factors are computed using the Random Draw model. The final BF is obtained by multiplying these BFs with those from the ASC family-based dataset, substituting any BF less than 1 with 1 to avoid down-weighting. We then convert the resulting BFs into FDR q-values and identify genes with q-values below 0.05. This procedure is repeated 100 times for each threshold level, and the average number of discoveries is reported. As in the null simulation, we use the result from a published dataset (Fu et al., 2022) as a proxy for the true risk gene status.

When the TADA model is applied to only ASC family-based data, the number of discoveries is 58, and when case data are considered within the TADA original model, TADA_*CC*, a total of 64 genes are discovered. TADA_*RD* provides a larger number of gene discoveries (Table 1). A threshold of 0.7 provides the best results in terms of achieving good power and yet controlling for errors.

### Integration of case data with the Random Draw model

We apply the proposed procedure to the ASC data (Figure 1) to identify risk genes associated with ASD. The ASC family-based dataset, which includes offspring data with parental information, is combined with datasets lacking parental information (ASC case-control). The family-based dataset is analyzed exclusively through the original TADA framework, which provides initial Bayes Factors for each gene. Case-only variants are evaluated by the classifier *ClassDn* to obtain de novo scores for them. Applying a threshold of 0.7 to de novo scores separates the variants into likely de novo and likely inherited sets using either of two types of *ClassDn* classifiers, RUSBoost and UnderBagging. Of the 13,486 PTV variants for case-only dataset, 2.2% (RUSBoost) and 3.7% (UnderBagging) of variants without parental information are classified as *de novo*. The Random Draw model then evaluates these pseudo-labeled data and generates the second BF using the sensitivity and specificity parameters learned from the ASC family-based dataset (Figure 3). After ensuring that each BF exceeds or is set to a minimum of 1, the two BFs are multiplied. For the final discoveries, the BFs are converted to FDR values (q-values) following the approach described in Li et al. (2010) (Section 2.3). This procedure requires an estimated proportion of risk genes, for which a value of 0.06 was used. The final discovery is confirmed by selecting genes with a q-value less than 0.05. This entire procedure is referred to as TADA_*RD*_.

The results highlight several features of our approach. First, *ClassDn* effectively separates different inheritance classes of variants in terms of the six covariates used in the classifier (Figure 2C). Likely de novo variants tend to have low LOEUF scores, high CCR, low frequency, low FDR TADA DD, low obs lof, and high exp lof, which is consistent with the properties of *de novo* variants observed in family-based data. For certain covariates, such as FDR TADA DD, the separation becomes more pronounced when pseudo-labels are used instead of actual *de novo* labels. In part, this occurs because only a subset of *de novo* variants is classified as likely de novo due to the conservative nature of the classifier when using a threshold of 0.7.

Next, TADA_*RD*_ significantly enhances the discovery of gene associated with ASD, identifying a total of 85 risk genes for RUSboost and 103 risk genes for underbagging. The traditional TADA framework, TADA_*CC*_ using only PTVs, identifies 64 genes. Because the final Bayes Factors are the product of two BFs, one derived from family data and the other from case-control data, for both TADA_*RD*_ and TADA_*CC*_, the contribution of each dataset can easily be decomposed by taking the logarithm of the final BFs (Figure 4; the result with RUSboost is shown). Under TADA_*CC*_, the contribution of case-control data is modest for most of the selected genes. Under TADA_*RD*_, however, the contribution is greatly increased for many genes. Case-only data contribute a substantial portion of the signal for many of the top-ranked genes, such as *POGZ, GRIN2B, KDM5B* and *MIB1*, which are considered high-confidence ASD genes in the literature (Fu et al., 2022).

**Fig. 4.**
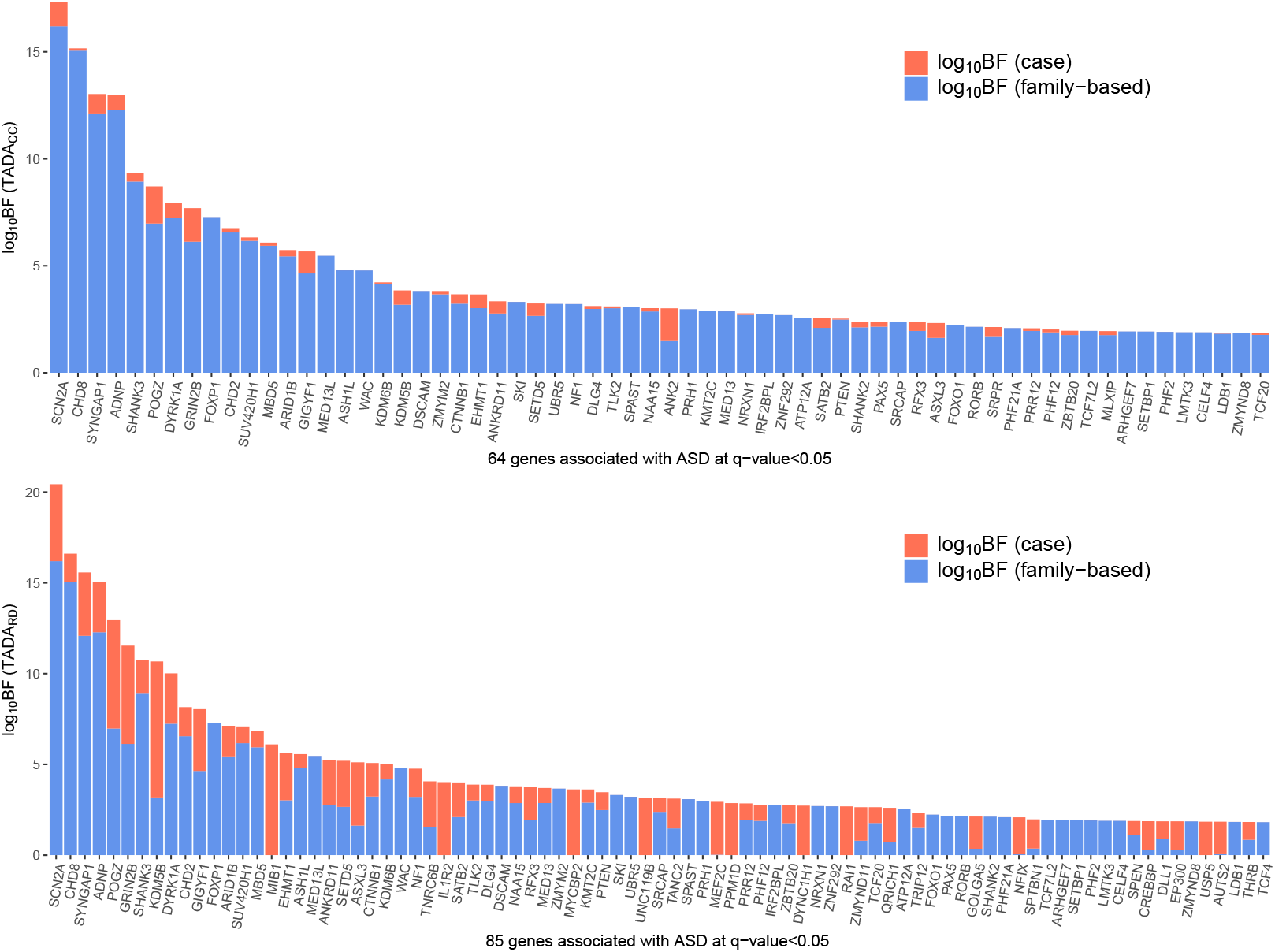
Integrating case data using the Random Draw model improves the association power among candidate genes. (Top) The evidence of ASD association contributed by each dataset for the 64 genes selected using the TADA_*CC*_ method with FDR ≤ 0.05. (Bottom) The evidence of ASD association contributed by each dataset for the 85 genes selected using the TADA_*RD*_ method with FDR ≤ 0.05. The result with RUSboost is shown. Most of the additional discoveries are validated by ongoing ASD cohort studies with larger datasets (unpublished yet).

Among the genes not selected by TADA_*CC*_, 23 are selected through both versions of *ClassDn* (Table 2). These selections are better understood by the number of likely de novo variants identified by *ClassDn*. Naturally, genes selected only by TADA_*RD*_ include at least one likely de novo variant in the case-only data. Some of these genes, such as *MEF2C* and *TNRC6B*, already exhibit moderate signal from TADA_*CC*_. However, for many other genes, case-only data serve as the primary source of the association signal: their q values are not even close to 0.05 in TADA_*CC*_, but are below 0.05 in TADA_*RD*_. This shows the critical boost in power of the new TADA model from inferred *de novo* variants and its capacity to elevate case-only data to one of the main sources of information for association. Two genes become insignificant when using TADA_*RD*_: *MLXIP* and *SRPR*.

**Table 2.**
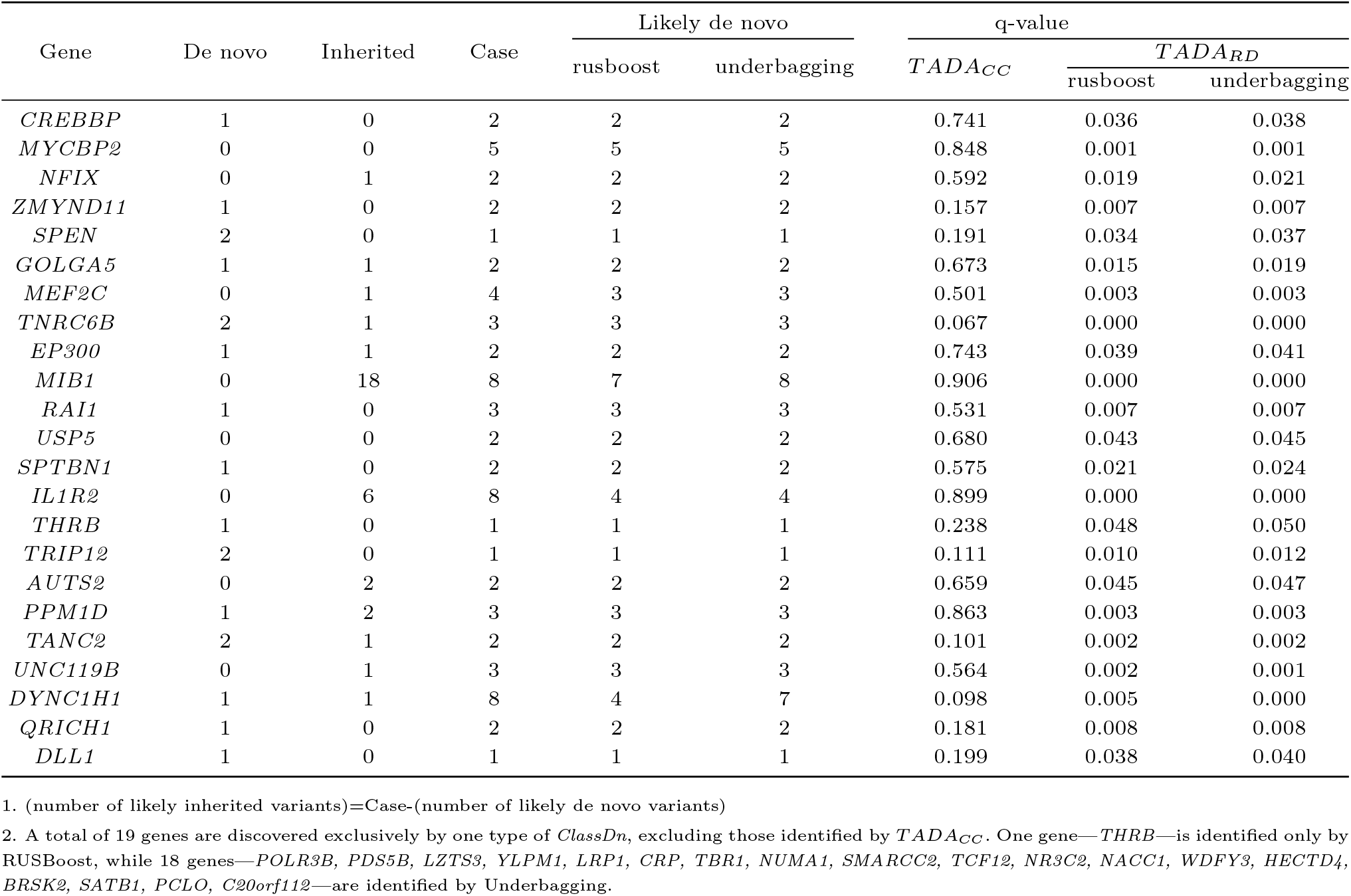
A list of 23 genes additionally discovered by *TADA*_*RD*_ compared to *TADA*_*CC*_, using a q-value cutoff of 0.05. Only genes commonly identified by both *ClassDn* methods (RUSBoost and Underbagging) are included.

## Conclusion and Discussion

Recent results show that the detection of *de novo* LoF mutations can be a powerful means of discovering novel risk genes for various developmental disorders, including ASD (Fu et al., 2022) and congenital heart disease (Jin et al., 2017; Watkins et al., 2019). Yet, *de novo* events are relatively rare, roughly one per exome. Thus, if we can increase the number of *de novo* mutations identified in probands, especially for probands without complementary parental information, we should be able to increase the power to identify risk genes. To address this challenge, we propose a novel framework that integrates both family-based and case-control data using a classification algorithm to probabilistically infer inheritance status of variation found in subjects who do not have complementary parental data. Then, by integrating a classifier trained on family-based data with a principled gene-level association model (A Random Draw model), we address the key challenge of missing inheritance labels in case-control or case-only datasets. Besides discovering additional risk genes, there is another key benefit of these methods. Using our classifier, likely *de novo* variants can be identified, which can be critical for clinical genetic evaluation when complete parental genotypes are unavailable.

Simulation studies demonstrate that this novel approach maintains control over false discoveries and can substantially enhance statistical power. Furthermore, when applied to exome sequencing data from ASD families and ASD case-control studies, the proposed method (TADA_*RD*) substantially increases the number of risk gene discovered compared to the conventional TADA model. This increased power emerges by using case data more effectively, specifically identifying likely *de novo* variation and interpreting the resulting *de novo* score using TADA_*RD*. Notably, most of the additional genes identified by TADA_*RD* are already supported by biological evidence or results of emerging large-cohort studies. One gene, *UNC119B*, is questionable because it has no other supporting evidence and may represent a false positive. Although this work focuses on protein-truncating variants, the framework is broadly applicable and can be extended to other variant classes and phenotypes. Our approach highlights the value of incorporating probabilistic annotations of *de novo* status into association models, and offers a scalable solution for maximizing information from incomplete but abundant sequencing datasets.

## Supporting information

Simulation of Denovo and Inherited PTV in Probands and Siblings

Result tables

## Acknowledgment

This work is supported in part by the National Institute of Mental Health (NIMH) [R01MH111658, R01MH129725] [R01MH128813, R01MH129724, U01MH111661], the Beatrice and Samuel A. Seaver Foundation, and New Faculty Startup Fund from Seoul National University.

## Supplementary Material A : Method Details

### Data preprocessing

*De novo data* We obtained all de novo and inherited variant data from Fu et al. (2022) for ASC and SPARK, separately. All de novo variants incorporated in this study were used directly as provided. Additional postprocessing filters were applied to inherited variants so that the final dataset was enriched for true associations. Specifically, we added an additional within-family segregation filter: variants were retained only if they were transmitted exclusively to affected children or exclusively to unaffected siblings, but not to both. This approach is based on the assumption that ultra-rare variants transmitted to both affected and unaffected siblings are unlikely to meaningfully contribute to autism risk. To further improve signal quality, we filtered inherited variants to those with allele frequency *leq* 0.00005 in the non-neuro subset of gnomAD v2.1.1 and *leq* 0.0001 in their respective internal datasets. Inherited PTVs were only retained if classified as high-confidence (HC) by the LOFTEE plugin for VEP and allowed only the “SINGLE EXON” flag among LOFTEE annotations. All variants were subsequently annotated with CCR scores (Havrilla et al., 2019), gene-level LOEUF metrics (Karczewski et al., 2020), expected and observed predicted loss-of-function variant counts (exp lof and obs lof, respectively), Developmental Delay FDR scores (Fu et al., 2022) and gnomad non-neuro AF, when available. Otherwise, variant frequencies were approximated within the appropriate dataset. Note that CCR coordinates were printed in bedfiles, which begin incrementing at 0. Therefore, we added 1 to all start and end coordinates to ensure compatibility with all VCF files.

*Case-Control data* We obtained case-control variant data from Fu et al. (2022). The case-control cohort contains 1,370 autism cases (*n*_*female*_ = 352, *n*_*male*_=1,018) and 4,249 unaffected controls (*n*_*female*_ = 2,023, *n*_*male*_ = 2,226). Variants were annotated with allele frequency data from gnomAD v2.1.1; when unavailable, allele frequencies were approximated using internal cohort-specific estimates. Consistent with the approach used for filtering and annotating de novo variation, case-control variants were also filtered using a non-neuro subset of gnomAD v2.1.1 allele frequency threshold of *leq* 0.001. As with the trio data, all variants were subsequently annotated with CCR scores (Havrilla et al., 2019), gene-level LOEUF metrics (Karczewski et al., 2020), expected and observed predicted loss-of-function variant counts (exp lof and obs lof), Developmental Delay FDR scores (Fu et al., 2022), and gnomAD non-neuro allele frequencies, when available.

### Details of the classifier training procedure

We use rare variant information from the SPARK family-based dataset, which includes data from 7,008 families. Each variant in this dataset is labeled with its inheritance class and accompanied by six offspring-level covariates that help distinguish between these classes. The fraction of *de novo* variants is approximately 0.0063 times that of inherited variants. In such scenarios, a naive application of a classifier tends to assign most variants to the inherited group—the majority class. While this approach may achieve a reasonable overall accuracy, it is not effective for the actual purpose of classification. This highlights the need for algorithms specifically designed to handle imbalanced data. Such algorithms typically rely on ensembling results from multiple learners, each trained on a subset of samples that are balanced through either oversampling the minority class or undersampling the majority class.

After an initial screening of various ensemble algorithms (results not shown), we selected two methods that use random undersampling of the majority class: RUSBoost (Seiffert et al., 2009) and Underbagging (Barandela et al., 2003). A key difference between these two algorithms is that RUSBoost iteratively selects training samples for each learner, assigning greater weight to samples that were misclassified by previous learners. In contrast, Underbagging builds learners in parallel, each trained on a randomly under-sampled, balanced subset of the data. The outputs of these algorithms are scores ranging from 0 to 1, where a higher score indicates that a variant is more likely to belong to the de novo class. A specific threshold can then be applied to these scores to complete the classification procedure. For example, applying a threshold of 0.7 means that a variant is classified as de novo only if its score exceeds 0.7.

RUSBoost and Underbagging require several tuning parameters. The imbalance ratio parameter refers to the intended imbalance ratio for each learner, defined as the ratio of majority instances to minority instances after class rebalancing. The size parameter specifies the number of learners. Additionally, a choice of algorithm is required to construct the learners. We use the random forest algorithm, which includes a parameter for selecting the number of trees. To select these parameters, we divided the dataset into five equally sized folds. Four folds were used to train the model, and the remaining fold was used for validation. The performance metric was constructed based on two scores: precision and recall, defined as:

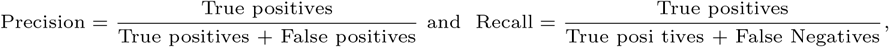

where we define *de novo* class as positive. Tested parameters include ir=1, 1.5, 2 ; es=10, 20, 30; ntree=20, 50, 100 (where appropriate). Note that the LOEUF composition of variants in each bin was roughly identical by design. The final decision on the tuning parameters was based on the area under the Precision-Recall (PR) curve, which evaluates the performance of binary classification algorithms across different threshold levels (Figure S1). The final AUC value is averaged over the results from five test folds.

**Fig. S1.**
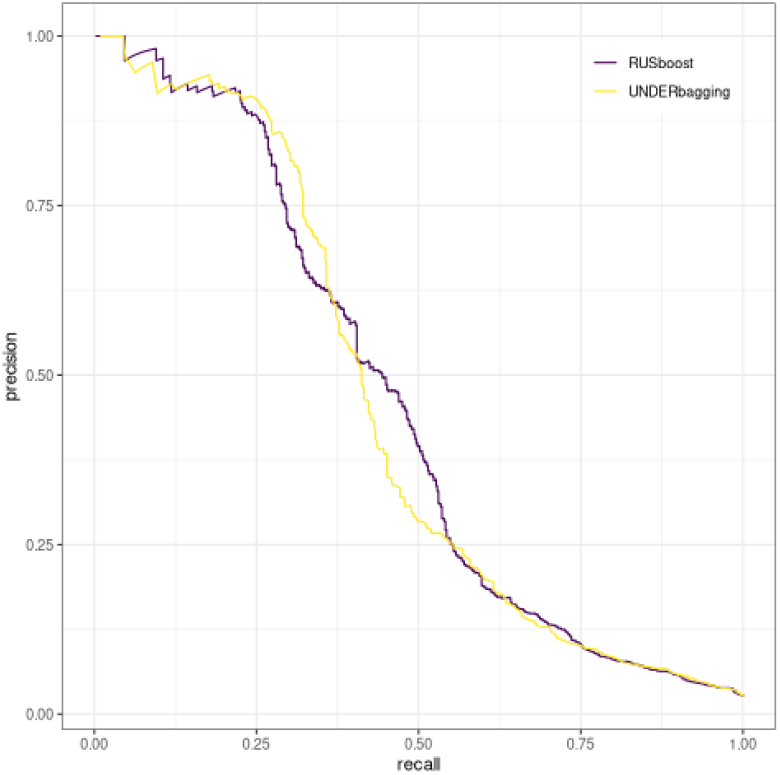
PR curve for parameters imbalance ratio=2, size=30, ntree=20 (fold 1)

The results show that the mean area under the PR curve does not vary greatly across different parameter choices, illustrating that the classifier is robust to the selection of parameters (Table S1). Considering both performance and computational complexity, we selected an imbalance ratio of 2, ensemble size of 30, and number of trees per learner as 20 for both RUSBoost-and Underbagging-based classifiers. The PR curves for this set of parameters are also largely similar between RUSBoost and Underbagging, suggesting that the overall performance of the two algorithms is comparable (Figure S1). In the main analysis, when we tested the trained classifiers on the ASC family-based data, we found that RUSBoost identified fewer *de novo* variants than Underbagging at the same threshold. This difference does not indicate that Underbagging outperforms RUSBoost; rather, it reflects that the same threshold corresponds to a lower point on the recall axis for the RUSBoost-based classifier.

**Table S1.** Mean PR AUC result for parameter selections.

| Model | Imbalance ratio | Size | ntree | Mean.PR.AUC |
| --- | --- | --- | --- | --- |
| rusboost | 1 | 10 | 20 | 0.404 |
| rusboost | 1 | 30 | 50 | 0.413 |
| rusboost | 1.5 | 10 | 20 | 0.427 |
| rusboost | 1.5 | 20 | 50 | 0.437 |
| rusboost | 2 | 20 | 50 | 0.451 |
| rusboost | 2 | 30 | 20 | 0.454 |
| rusboost | 2 | 30 | 100 | 0.453 |
| underbagging | 1 | 10 | 20 | 0.411 |
| underbagging | 1 | 30 | 50 | 0.426 |
| underbagging | 1.5 | 10 | 20 | 0.434 |
| underbagging | 1.5 | 20 | 50 | 0.443 |
| underbagging | 2 | 20 | 50 | 0.454 |
| underbagging | 2 | 30 | 20 | 0.457 |
| underbagging | 2 | 30 | 100 | 0.459 |

### Details of the Random Draw Model

The random draw model first constructs the likelihood of having observed values of likely de novo and likely inherited variants under each risk gene and non-risk gene scenario. It then calculates a Bayes factor, defined as the ratio of this two likelihoods, as the evidence supporting the gene being a risk gene.

#### Background of the random draw model

The *TADA* models is a well-established framework to model the number of *de novo* and inherited variants based on genetic parameters under both risk gene and non-risk gene scenarios. Both *de novo* and inherited variants contribute to the evidence supporting the gene being risky. A likelihood ratio, based on the observed number of variants, combined with estimated genetic parameters, provides a risk score for a specific gene. While both *de novo* and inherited variants contribute to the evidence supporting the gene being risky, results from Fu et al. (2022) show that while the gene risk score is positively correlated with the number of *de novo* variants, it is negatively correlated with the number of inherited variants (Figure S2). Most risk genes have a very limited number of inherited variants, while they have a relatively many number of *de novo* variants.

**Fig. S2.**
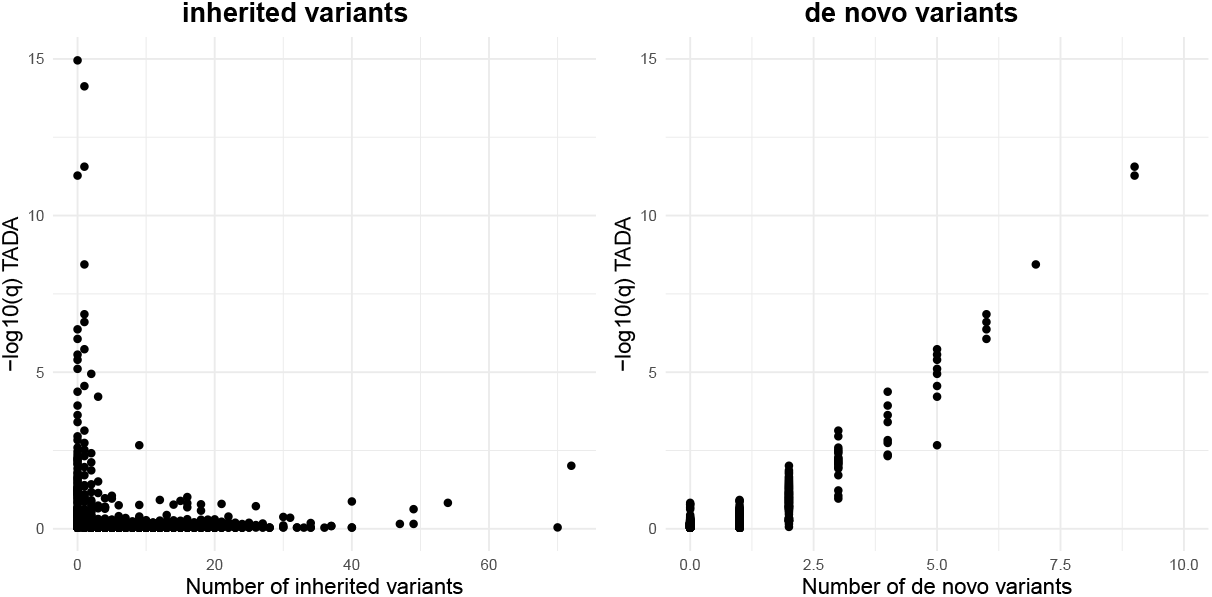
Marginal relationships between gene risk scores and the number of variants per gene: the number of inherited variants (left) and the number of de novo variants (right). Gene risk scores are represented as -log(q-value) computed in Fu et al. (2022).

The random draw approach models this marginal relationship rather than establishing structural equations for the number of variants. Specifically, it assumes that each variant is a randomly drawn observation from a mixed pool of *de novo* and inherited variants and estimates the likelihood of the observed composition of *de novo* and inherited variants. Generally, the probability of a random variant being *de novo* is higher for risk genes. Therefore, given a fixed total number of variants, the more *de novo* variants there are, the more likely the gene is to be a risk gene.

With this approach, we can effectively deal with the uncertainty of the inheritance class involved in the case data, as explained in the subsection below. Additionally, even though this approach takes a different perspective to model the observed numbers of variants, it remains highly consistent to the original TADA model when there is no uncertainty regarding the inheritance class (see Figure S3).

**Fig. S3.**
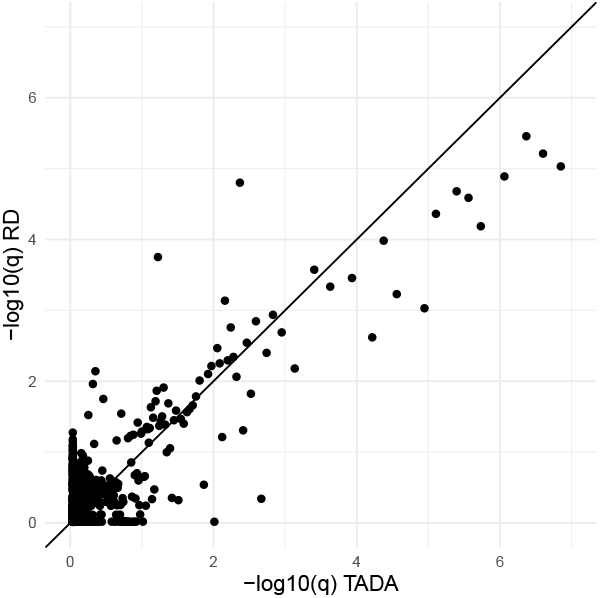
Comparison of the Random Draw and the TADA model in family-based data.

#### Formulation

Let *D*_*i*_ denote the risk status of the gene *i*, where *D*_*i*_ = 1 if the *i*th gene is a risk gene and *D*_*i*_ = 0 otherwise. Let 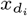 and 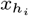 denote the number of likely de novo and likely inherited variants, respectively, oberved in gene *i*. Also, let 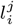 denote the likely denovo status of the single variant *j* of gene *i*; 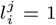 if it is likely de novo and 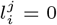 if it is likely inherited. Then the likelihood of observing 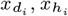 under risk and non-risk gene scenarios are formulated as follows:

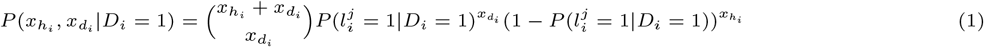

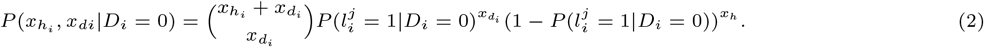

The quantities 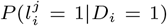 and 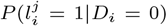 denotes the probability that randomly selected variant is likely de novo for risk genes and non-risk genes, respectively. Let 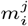 be the indicator of variants being *de novo*, where 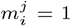 if *de novo* and 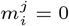 otherwise. Then we have

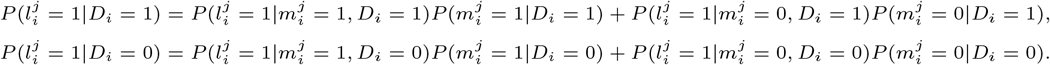

Here we assume that the probability that a *de novo* variant is observed as likely de novo and a inherited variant is observed as likely inherited is indifferent between risk and nonrisk genes, that is :

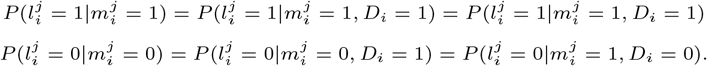

Then, we can compute the probability that some randomly drawn variant from gene *i* is likely denovo:

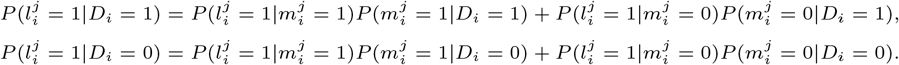

Two performance parameters 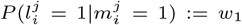 and 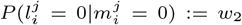 denotes the sensitivity and specificity of the classifier, respectively. They can be estimated empirically in a test sample (ASC family-based data in our application). Also, the quantities 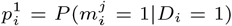 and 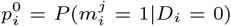 are genetic parameters representing the probabilities that a variant drawn from gene *i* is a de novo variant, under the risk-gene and non-risk-gene scenarios, respectively. The parameter 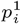 is expected to be higher than 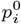. For given 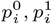, we have

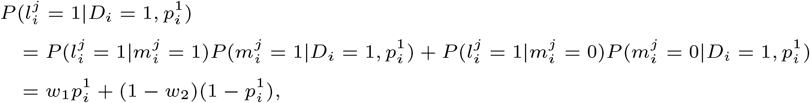

and

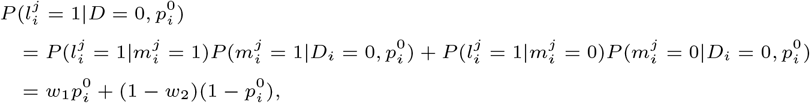

In our model, the parameters 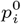 and 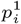 follows some nondegenerate distributions under each risk and nonrisk scenarios: 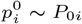 and 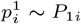. To estimate these distributions, we specify the prior distributions *p*_0*i*_ ∼ *Beta*(*α*_0*i*_, *β*_0*i*_) and *p*_1*i*_ ∼ *Beta*(*α*_1*i*_, *β*_1*i*_) and estimate the gene-specific hyperparameters as follows. We group the genes based on their pre-risk status, which are determined by whether the q-values from the TADA model applied to only family-based data are above or below 0.05 (Fu et al., 2022). Then, for each pre-risk and pre-nonrisk gene sets, we fit a logistic regression model for the *de novo* ratio against *log*_10_(mutation rate) for each gene, and use the predicted values to represent the mean values *µ*_0*i*_ or *µ*_1*i*_ of the distributions *P*_0*i*_ or *P*_1*i*_. The variance of *P*_0*i*_ or *P*_1*i*_ are estimated through a jackknife approach, separately, under the assumption that the variance are identical among genes within risk or nonrisk sets. To calculate such variances, we exclude one gene at a time and compute a mean ratio as the total number of *de novo* variants divided by the total number of variants in each pre-risk/pre-nonrisk group. Let 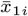 be the estimated mean of the *de novo* ratio calculated for genes in each group excluding the *i*th gene, and let ℐ_risk_ and ℐ_nonrisk_ are index sets for the pre-risk and pre-nonrisk genes. Then, the jackknife estimator for the variance is provided by

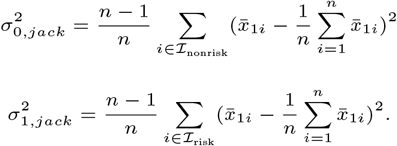

Then we convert gene-specific means and variance using formulas for beta distribution parameters;

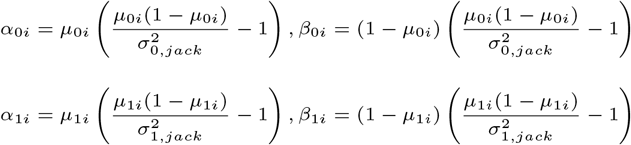

Finally, the evidence of *D* = 1 against *D* = 0 can be calculated as a Bayes Factor, which accounts for uncertainty in the parameters (*p*_0*i*_, *p*_1*i*_) using gene-specific prior distributions *P*_0*i*_ and *P*_1*i*_:

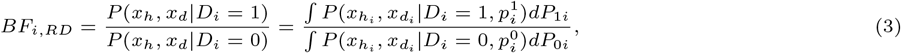

where the likelihoods are integrated over the distribution of 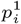 and 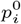.

#### Robust check for the random draw model

To further validate the model, we compare the result from the random draw model and the original TADA model. In our main analysis, the random draw model was only applied to a case data, while the evidence from the family-based data is still collected through the family-based component of the *TADA*_*CC*_ data. That is, in our main implementation, the only difference between *TADA*_*RD*_ and *TADA*_*CC*_ model is the treatment of the case-control data part. However, the random draw model can be still applied to a family-based data by simply setting *w*_1_ = *w*_2_ = 1. Since the TADA model has been validated in numerous previous studies for its validity and effectiveness, the comparison between the TADA model and the random draw model for the family-based data will serve as a tool to assess the robustness of our random draw model. A high correlation between two different approaches supports the validity of both methods.

