## Supplementary material for "A framework to infer *de novo* exonic variants when parental genotypes are missing enhances association studies of autism": Simulation of Denovo and Inherited PTV in Probands and Siblings

**Supplementary Material B : Simulation of Denovo and Inherited PTV in Probands and Siblings**

This document illustrates the procedure for generating a realistic simulation of de novo and inherited PTVs in probands and siblings. The results were used in the manuscript to perform a null simulation analysis and to generate a realistic case dataset for demonstrating the procedure in the accompanying code package.

1. **Simulation of Denovo and Inherited PTV in Siblings of Cases**

To create a life-like simulation, information was taken from various sources and combined into a master set of protein truncating variants (PTV). This set was subsequently used to simulate denovo PTV-variants based on the gene specific mutation rates for PTVs and inherited PTVs using the PTV-variant specific gnomAD non-neurological samples.

**Sources of Information and Data Preparation**

***Gene and transcript information***

A list of 80,950 transcripts representing 19,724 genes was obtained from the LOUEF file (supplementary_dataset_11_full_constraint_metrics.tsv.gz; downloaded: 2023-11-21). After removing 1776 transcripts for which no LOEUF score was available, 79,174 (19,233 genes) remained. Subsequently, biomaRt (accessed: 2025-05-12) was used to determine: 1) whether the transcript was still supported; 2) the genomic boundaries of the transcript based on build38 (HG38); and 3) whether the transcript is the canonical transcript for the gene. The biomaRt based filter reduce the number of transcripts to 71,876 (19,004 genes).

***PTV information***

The next step was to process the variant information from gnomAD file with variant information (downloaded: 2025-02-27). The chromosome files were screened for PTV variants using the following criteria: 1) PASS field equals PASS; 2) IMPACT equals HIGH; 3) BIOTYPE equals protein_coding; 4) Exonic = T; 5) minor allele frequency based on non-neuro samples < 0.001; and 6) non-missing consequence. This screen of the gnomAD data yielded a total 323,264 PTV variants in 17,376 transcripts representing 16,372 genes. The gnomAD information supplied location information for the variants based on build38 (HG38). PTV-variants were of the types frameshift, stop gained, stop lost, start lost, splice acceptor, and splice donor.

***Combining gene/transcript and PTV information***

The number of overlapping transcripts between the gene information and the variant information was 15,278. This set represents 14,557 unique genes and a total of 282,766 ptv variants.

For 117 transcripts the Ensembl gene id reported in the LOUEF file did not match the id reported by biomaRt. Removal of these transcripts resulted in a set of 15,161 transcripts, 14,461 genes, and 281,049 PTV variants.

Based on the biomaRt information 10,146 of the transcripts were identified as canonical. An additional 1464 transcripts were identified as canonical based on the LOEUF information. For each gene a single transcript was chosen to represent it based on canonical status (TRUE over FALSE). Ties were broken based on the coding sequence length of the transcript (longer over shorter). This led to a list of 14,461 genes and transcripts and 273,042 variants.

***PTV mutation rate***

Gene specific mutation rates for PTV based on ASC samples were obtained from the Fu paper Supplementary Table 5 (downloaded: 2022-09-28). Matches with our data could be made for 14,451 genes. For the remaining 10 genes, the mutation rates reported in the LOUEF data were used.

***Conserved coding region information***

Information on 8,188,409 conserved coding regions (CCR) was obtained from the file ccrs.autosomes.v2.20180420.bed.gz (downloaded: 2025-02-25). After removing genes for without a matching Ensembl gene id, 8,107,516 regions in 17,105 genes remained. Since the CCR location was provided based on build37 (HG19), the liftOver script (downloaded: 2021-11-21) was used to perform a lift-over to build38 (HG38). Proper lift-over of genomic coordinates to build38 (HG38) could be obtained for 8,106,743 CCR regions.

***Combining source of information***

Some final quality control of the data removed 473 variants that were not within the genomic location boundaries of a canonical transcript. This resulted in a useable variant dataset of 272,929 variants representing 14,447 genes/transcripts. The CCR information is subsequently added to the PTV based on its location. In case of a PTV mapping to multiple CCR regions, the maximum ccr_per is used. CCR values could be assigned to 244,734 PTV variants. The remaining 25,195 received a missing value, although one could argue that a ccr_per=0 should be used for these cases.

**Simulation**

The purpose of the simulation is to generate a dataset of rare denovo and inherited PTV variants for siblings of cases that is similar to what is observed for the ASC. The ASC data is characterized as being a set of 5374 PTV variants occurring in 3768 genes. There are 5121 variants that are identified as being inherited, 189 that are denovo, and 4 that are observed as being inherited and denovo. Of the variants, 65 have ccr_per > 90%, 883 have a ccr_per between 1 and 90, 2786 have ccr_per = 0, and the remaining 1571 variants did not occur in a CCR region. The variants were divided into deciles based on the reported gnomAD allele frequency. Cut-offs for the quantiles are in Table 1. This information will be used to down sample the simulated dataset to the required number of variants

**Table 1.** Minor allele frequency cut-offs for the gnomAD frequency deciles observed in the ASC data. Variants without a reported gnomAD frequency were excluded from this summary.

| Decile | Minor allele frequency cut-off |
| --- | --- |
| 10% | 4.88204e-06 |
| 20% | 9.70400e-06 |
| 30% | 1.92230e-05 |
| 40% | 2.91664e-05 |
| 50% | 5.10495e-05 |
| 60% | 8.36136e-05 |
| 70% | 1.44275e-04 |
| 80% | 2.49586e-04 |
| 90% | 4.46125e-04 |
| 100% | 1.00000e-03 |

The simulation starts with the user providing the number of samples to use to determine denovo variants (n.dn) and the number of samples to use for generating inherited variants (n.inh). Next the user can provide the detection rate for inherited variants. While testing it appears that it is best to leave this at 1. Final input is the seed. Using NULL will allow for a random seed.

The code relies on two external files which are supplied with the code:

1) gene-hg38-df-with-ccr_2025-03-25.txt – gene level data
2) ptv-hg38-df-with-ccr_2025-03-25.txt – variant level data

***Algorithm for Denovo Variants***

The number of carriers of denovo PTV variants for a gene and the PTV variants that they carry is determined using the following algorithm:

1) randomly determine the number of carriers, m, using a Poisson with λ = 2×n.dn×ptv mutation rate of the gene.
2) randomly sample with replacement m PTV variants from the community of PTV variants of the gene with replacement.
3) Tabulate which and how often the PTV variants were selected.

***Simulating Inherited Variants***

A rare inherited PTV is defined as a variant that occurs in the child and in at least one of the parents. The PTV cannot occur as a homozygote in any of the members of the n.inh trios.

The number of carriers of an inherited PTV, m, were simulated using the following algorithm.

1) Simulate genotypes for the n.inh fathers and n.inh mothers using a binomial distribution with size = 2 and p = gnomAD minor allele frequency.
2) Exit with m = NA, if there is at least one father or mother with a minor allele homozygous
3) Generate the genotype of the children by randomly choosing which allele is being transmitted from the father and the mother using a Bernoulli distribution with p = 0.5.
4) Exit with m = NA when there is at least one child with a minor allele homozygous genotype.
5) Return m, the number of children that are heterozygous for the minor allele.

(Comment: the code is set up to allow for variant detection rates < 1. This is to mimic the situation in which sequencing depth is likely not to detect all occurrences of the variant. For this application, this is not critical).

***Down Sampling Variants***

Using the two algorithms using the 272,929 PTV in 14,447 genes, assuming n.dn = n.inh = 2500, yields ~185 denovo PTV and ~15.6K inherited PTV occurring in ~7300 genes. While the number of denovo PTV is similar to what is seen in the ASC data, the number of simulated inherited PTV far exceeds the number observed in the ASC. Explanations for this could be a deeper sequencing depth in gnomAD versus ASC, a more restrictive calling algorithm in the ASC, different ancestry composition in the two datasets, among others.

A closer inspection of the simulated result revealed that among the 15.6K there are approximately ~600 with 1% ≤ ccr_per < 90%, and ~50 with cct_per ≥ 90%. These are about half of what we see in the ASC data.

***Algorithm for down sampling:***

1) Randomly draw a number between 5000 and 5500 to determine the number of PTV required (N). The number of PTV in each minor allele decile is then N/10.
2) Accept all denovo PTV, determine their decile bins.
3) Accept the inherited PTV with ccr_per ≥ 1, determine their decile bins.
4) Adjust the number of PTV need for each decile bin by the number of PTV assigned to the bin in step 2) and 3).
5) The sum of the adjusted number is the total number of inherited PTV with ccr_per < 1 or missing needed to fill up the bins (k).
6) Determine the decile bin for inherited PTV with ccr_per < 1 or missing.
7) Calculate the ratio of the number of PTV needed from each bin (step 4) divided by the number available (step 6)
8) Randomly draw k PTV, without replacement, from the inherited PTV with ccr_per < 1 or missing, using the weights from step 7)

Using this algorithm, results in a dataset with N variants, and in my example, ~3700 genes.

2. **Simulation of Denovo and Inherited PTV in Siblings of Cases**

This simulation follows the general simulation of the denovo and inherited PTV in siblings of cases. Instead of relying on random occurrences of these events, the counts in the ASC data for the 185 significant genes in the Fu paper are used to develop a distribution of the number of denovo and inherited events in the significant genes.

**Determining the distribution of denovo and inherited events for genes with a signal**

Table 1 shows the distribution of the observed number of genes out of 185 with their number of denovo and inherited events. Observed entries in this Table were then used to fit a model log$\left( n_{gene} \right)\sim n_{event}+n_{event}^{2}$. The R^2^ for the denovo and inherited data were 0.92 and 0.90, respectively. Observed counts with value NA were not used in developing the prediction model. The regression results were used to determine the expected number and fraction of genes in the event counts 0 through 12. The fraction will be used in weighted sampling scheme to assign number of events to signal genes. Of note is that one of the genes had an inherited event count of 39. For the regression, I decided to replace this count by 12.

**Table 1.** Distribution of the number of denovo and inherited events in the ASC data for the 185 significant genes from Fu.

|  | denovo | | | inherited | | |
| --- | --- | --- | --- | --- | --- | --- |
| # events | observed | predicted | fraction | observed | predicted | fraction |
| 0^1^ | 86 | 84.6 | 0.4434 | 112 | 90.2 | 0.5307 |
| 1 | 30 | 42.9 | 0.2248 | 35 | 36.8 | 0,2166 |
| 2 | 27 | 23.1 | 0.1211 | 17 | 16.7 | 0.0981 |
| 3 | 15 | 13.2 | 0.0692 | 8 | 8.4 | 0.0492 |
| 4 | 10 | 8.0 | 0.0420 | 3 | 4.7 | 0.0274 |
| 5 | 4 | 5.2 | 0.0271 | 3 | 2.9 | 0.0169 |
| 6 | 7 | 3.5 | 0.0186 | 2 | 2.0 | 0.0116 |
| 7 | 2 | 2.6 | 0.0135 | NA | 1.5 | 0.0088 |
| 8 | NA | 2.0 | 0.0104 | NA | 1.3 | 0.0074 |
| 9 | 1 | 1.6 | 0.0086 | NA | 1.2 | 0.0070 |
| 10 | NA | 1.4 | 0.0075 | 2 | 1.2 | 0.0072 |
| 11 | 1 | 1.3 | 0.0069 | 2 | 1.4 | 0.0083 |
| 12 | 2 | 1.3 | 0.0068 | NA | 1.8 | 0.0106 |
| 39^2^ |  |  |  | 1 |  |  |

^1)^ An observed count of 0 occurs because the results from Fu were based on two datasets, ASC and SPARK.

^2)^ For the prediction of the inherited, I choose to use a count of 12 instead of 39.

**Gene specific mutation rate**

In the ASC data, the approximately 8000 probands and 2500 siblings have a total of 1171 and 201 denovo PTV mutations. This suggest that the mutation rate in probands is 1.8 times larger in probands than siblings. For the simulations of denovo ptv events in probands the gene specific mutation rates that are used for siblings are therefore multiplied by 1.8 before using them in the proband simulation

**Simulation approach**

From the complete set of genes, randomly select N genes to be the genes with signal. For each of these genes, randomly assign the number of denovo (dn_gene_) and inherited (inh_gene_) events between 0 and 12, using the fractions from Table 1 as weights.

**Denovo simulation**

1) Randomly generate the number of denovo events for all genes based on the gene specific mutation rate and the number of probands in the simulation.
2) Retain the set with 1 or more denovo events
3) Remove the signal genes from this set
4) Add the signal genes with their dn_gene_
5) Randomly select, with replacement, the ptv variants in a gene to represent the denovo counts for each gene.

**Inherited Simulation**

1) Simulate inherited variants as for the sibling simulation. This includes the pruning to balance the variants across the CCR and MAF categories.
2) Remove variants found in the set of signal genes
3) For each of the signal genes with inh_gene_ > 0
   a. Generate the number of inherited events for each PTV in the gene using the approach used in the siblings.
   b. If n_inh_ is not met, repeat a) and add to the previous number of events for the gene
4) Add the results from 3) to the ones from 2)
